# IL-15 enhances functional properties and responses of cytotoxic CD4^+^CD28^−^ T cells expanded in Systemic lupus erythematosus

**DOI:** 10.1101/2023.01.04.522682

**Authors:** Tingting Wang, Laiyou Wei, Shuihui Meng, Wencong Song, Yulan Chen, Heng Li, Qianqian Zhao, Zhenyou Jiang, Dongzhou Liu, Huan Ren, Xiaoping Hong

**Affiliations:** Department of Rheumatology and Immunology, The Second Clinical Medical College, Jinan University (Shenzhen People’s Hospital), Shenzhen, 518020, China; Integrated Chinese and Western Medicine Postdoctoral Research Station, Jinan University, Guangzhou, 510632, China; School of Medicine, Southern University of Science and Technology, Shenzhen, 518055, China; Shenzhen People’s Hospital, The Frist Affiliated Hospital of Southern University of Science and Technology, Shenzhen, 518055, China; Department of Microbiology and Immunology, College of Basic Medicine and Public Hygiene, Jinan University, Guangzhou, 510632, China

**Keywords:** Interleukin-15, Cytotoxic CD4^+^CD28^−^ T cells, Inflammation, Cytolytic function, Renal damage

## Abstract

Systemic lupus erythematosus (SLE) is an autoimmune disorder that results in an attack by body’s immune system of its own tissues, causing chronic inflammation and tissue damage. T cells play a key role in the pathogenesis of SLE, as they secrete pro-inflammatory cytokines as well as mediate direct effects on target tissues. Recently, CD4-positive T cells with cytotoxic potential were showed to be involved in autoimmune disease progression and tissue damage. However, whether this cell type expands and plays effector functions in SLE patients remain to be elucidated. Analyzing single-cell RNA sequencing (scRNA-Seq) data and flow cytometry data, we find that cytotoxic CD4^+^CD28^−^ T cells are present in SLE patients. We also show that these cells expand most prominently in patients affected by lupus nephritis, and they exhibit cytotoxic activity against human renal glomerular endothelial cells in vitro. In addition, our study suggests that Interleukin-15 (IL-15) promotes the expansion, proliferation and cytotoxic function of CD4^+^CD28^−^ T cells in SLE patients. Tofacitinib, a selective JAK3 inhibitor, inhibits the effect of IL-15 on CD4^+^CD28^−^ T cells. Together, our study clearly demonstrated that CD4^+^ CD28^−^ T cells characterized by proinflammatory properties and cytolytic function expand in SLE patients. The pathogenic potential of these CD4^+^ CD28^−^ T cells is driven by IL-15/IL-15R/JAK3/STAT5 signaling pathway, which may open new avenues for therapeutic intervention to prevent progression of SLE patients.

## 1. Introduction

Systemic lupus erythematosus (SLE) is a complex autoimmune disorder characterized by an aberrant cytokine milieu as well as by overproduction of pathogenic autoantibodies[1, 2]. SLE constitutes a prototypic systemic disease that affects nearly all organs and tissues, and the kidney is one of the common targets of damage[3, 4]. Various immunological abnormalities related to the disease have been identified and overriding abnormalities are hyperactivity of B and T lymphocytes[5, 6]. B cells, as antibody-producing cells, have been studied extensively. T cells are now recognized as the key mediators in pathogenesis of both SLE and Lupus nephritis[7–10]. T cells represent the majority of renal-infiltrating immune cells[11, 12] and are critically involved in the development of organ inflammatory and damages. CD4^+^ T cells, including T helper type 1 (Th1), T helper type 2 (Th2), T helper type 17 (Th17), Regulatory T cells (Tregs), follicular helper T (Tfh) cells, as helpers and regulators of immune responses, have been reported to initiate and perpetuate inflammation and play critical role in the immune pathogenesis of SLE and Lupus nephritis[13–17].

CD4^+^ T cells with cytotoxic activity are a specialized CD4^+^ T subset, and the existence of these cells has been reported in several autoimmune and chronic inflammatory disease, including rheumatoid arthritis (RA)[18, 19], multiple sclerosis (MS)[20], acute coronary syndrome (ACS)[21, 22] and IgG4-related disease (IgG4-RD)[23, 24]. CD4^+^ cytotoxic T cells are characterized by a lack of expression of the surface co-stimulatory molecule CD28. Furthermore, these cells were previously shown to express cytotoxic-related genes granzymes and perforin and share many phenotypic features with CD8^+^ T cells and Natural Killer (NK) cells[25, 26]. Upon activation, CD4^+^ cytotoxic T cells release perforin, granzymes as well as inflammation cytokines. Secretion of this arsenal of chemical weaponry promotes the death of target cells in a number of ways[27]. Together, these functions of CD4^+^ cytotoxic T cells support the idea that the expansion of these cells plays a pathogenic role in the development of disease. Thus, understanding the complex functions of CD4^+^ cytotoxic T cells should facilitate therapeutic intervention to prevent disease development. However, the highly variable combinations of surface biomarkers used to defined CD4^+^ cytotoxic T cells, including cytotoxic and regulatory T cell molecule (CRTAM)[28], CX3C motif chemokine receptor 1 (CX3CR1)[20], natural killer group 2D (NKG2D)[29, 30], and SLAM family member 7 (SLAMF7)[24, 31], rendering functional comparisons challenging. In SLE, previous studies have shown that CD4^+^CD28^−^ T cells with the capacity of pro-inflammatory secretion and higher proliferative activity, were expanded in peripheral blood of patients with moderately active SLE. The percentage of CD4^+^CD28^−^ T cell positively correlated with the Systemic Lupus International Collaborating Clinics/ ACR Damage Index (SDI)[32, 33]. CD4^+^CD28^−^ T cells have previously been reported to predict the occurrence of new lung damage in SLE patients[34]. However, the phenotypic and functional properties and the possible pathogenic involvement of the expanded CD4^+^CD28^−^ T cells in SLE remain to be elucidated.

Here, we set out to characterize the phenotype and function of cytotoxic CD4^+^CD28^−^ T cells in SLE patients using single-cell transcriptomic data and flow cytometry. We investigated the effects of IL-15 on CD4^+^CD28^−^ T cells in SLE patients and inquired into the underlying molecular mechanisms.

## 2. Materials and Methods

### 2.1. Study subjects

A total of 149 patients with SLE were enrolled from the Department of Rheumatology and Immunology, Shenzhen People’s Hospital, including 80 patients with nephritis (LN group) and 74 patients without nephritis (NLN group). 39 healthy individuals (age and gender matched) were recruited as controls (Supplementary Table 1). Patients with other autoimmune diseases or infectious diseases or over 70 years old were excluded from this study. Blood samples were obtained from all patients and healthy controls. All patients fulfilled the Systemic Lupus International Collaborating Clinics (SLICC) classification criteria for SLE[35]. The Systemic Lupus Erythematosus Disease Activity Index 2000 (SLEDAI-2K) score for patients is calculated using the SLEDAI-2K score calculator according to both clinical and laboratory parameters (https://qxmd.com/calculate/calculator_335/sledai-2k)[36]. The SDI[37] was also assessed. The diagnosis of LN was confirmed by renal biopsy as per the International Society of Nephrology and Renal pathology Society (ISN/RPS) criteria[38]. Demographic data, clinical data and laboratory data were collected from hospital records or by questionnaire and reviewed by experienced physicians. This study was approved by the Ethics Committee of the Shenzhen People’s Hospital, China (LL-KY-2021703), and informed consent was obtained from all subjects according to the Declaration of Helsinki.

### 2.2. Single-cell Analysis

Raw sequencing data was processed with the Cell Range Pipeline (Version3.1.0), and the data was filtered, scaled, quantified, and finally the gene expression matrix of each cell was obtained. In brief, the sequencing data in FASTQ files was aligned to the GRCh38 (hg38) genome reference and transcriptome with cell range (Version3.1.0) pipeline and generated gene–cell expression matrix. The data were filtered with the following parameters: genes detected in >5 cells, and cells with 200~2000 detected genes and UMI with 1000~10000 and <15% mitochondrial genes. A total of 133,804 cells were obtained from 21 samples for further analysis. Next, FindIntegrationAnchors and IntegrateData function were used to integrate all sample datasets and to scale all expression data with ScaleData function. Principal component analysis (PCA) of the genes from these selected cells were performed, and 40 PCs were selected to identify clusters using the FindClusters function (40 PCA components with resolution 0.5 – 1.5, leading to 10-24 clusters, and a resolution of 0.4). Lastly, the Run TSNE function was used to perform t-Distributed Stochastic Neighbor Embedding (t-SNE) analysis, and the cells were annotated according to the expression of canonical lineage marker. Subsequently, the data of subsets (all CD4^+^ T cells in T cells) were analyzed. A total of 9462 cells were obtained for subsequent subgroup analysis. 10 PCA components with resolution 0.1 – 0.5, leading to 3-15 clusters, and a resolution of 0.1 were used for clustering. Finally, cells were annotated based on known and high scoring genes.

### 2.3. Flow cytometry

Blood samples were collected in EDTA anticoagulant tube, and 100 μl blood samples were transferred to a new tube for flow cytometry analysis. Peripheral blood mononuclear cells (PBMCs) were isolated using density gradient centrifugation. The frequency of CD4^+^CD28^−^ T cells and their phenotypical characterization were determined by staining blood with antibodies, followed by erythrocyte lysis with Lyse/Fix buffer (BD Biosciences). The percentage of circulating CD4^+^CD28^−^ T cells was calculated as the percentage of CD4^+^ T cells. For intracellular staining, including staining for cytotoxic molecules (granzyme B (GZMB), granzyme A (GZMA), perforin (PRF1)), transcription factor (T-bet, Runx3) and cytokines (Interferon gamma (IFN-γ), tumor necrosis factor alpha (TNF-α)), PBMCs or cells from whole blood after surface marker staining and erythrocyte lysis, was fixed, permeabilized and stained with antibodies using a Fixation/ Permeabilization Solution Kit (BD Biosciences) according to the manufacturer’s instructions. For assessing the Phosphorylation state of signal transducer and activator of transcription 5 (STAT5), CD4^+^CD28^−^ T cells were isolated and then cultured alone or treated with IL-15. Phosphorylated STAT5 and extracellular-signal regulated kinase (ERK) were assessed via the BD Phosflow method using STAT5 antibody, ERK1/2 antibody, BD Phosflow Lyse/Fix Buffer, and BD Perm Buffer III (all BD Biosciences) as per the manufacturer’s instructions.

Antibodies targeting the following molecules were used in this study: CD3 (clone:HIT3a; PerCP/Cy5.5 BioLegend, clone:UCHT1; Brilliant Violet421 BioLegend), CD4 (clone:A161A1; APC-Cy7, PE/Cyanine7 BioLegend), CD8 (clone:SK1; APC-Cy7 Biolegend and BD Biosciences), CD28 (clone:CD28.2; FITC, PE-Cy7 BioLegend), granzyme B (clone:QA18A28; PE Biolegend, clone:QA16A02; Alexa Fluor 647 Biolegend), granzyme A (clone:CB9; PE Biolegend), perforin (clone:dG9; PE/Cyanine7 BioLegend), CX3CR1 (clone:2A9-1; APC eBioscience), NKG2D (clone:1D11; APC BD Biosciences), T-bet (clone:4B10; PE/Cyanine7 BioLegend), Runx3 (clone:R3-5G4; PE BD Biosciences), TNF-α (clone:MAb11; APC BioLegend), IFN-γ (clone:4S.B3; PE BioLegend and BD Biosciences, PE/Cyanine7 BD Biosciences), STAT5 (pY694; PE BD Biosciences), ERK1/2 (pT202/pY204; PE/Cyanine7 BD Biosciences). Data were acquired using FCAS Canto II (BD Biosciences) and analyzed with FlowJo software.

### 2.4. Intracellular cytokine staining

Intracellular cytokine assay was performed to assess cytokine levels. Briefly, PBMCs were stimulated with a combination of phorbol 12-myristate 12-acetate (PMA) and ionomycin (both from Sigma-Aldrich) for 4 hours. Three hours prior to analysis, Brefeldin A (Sigma-Aldrich) was added. PBMCs were stimulated in standard culture medium without PMA and ionomycin (Ion) as a control. To analyze IFN-γ and TNF-α producing cells, the cell surface of PBMCs was stained with antibodies (anti-human CD3, anti-human CD4, anti-human CD28). Next, cells were fixed, permeabilized and stained with antibodies (anti-human IFN-γ and anti-human TNF-α).

### 2.5. Degranulation assay

PBMCs were isolated from 10 ml fresh peripheral blood samples by Ficoll-Paque (GE Healthcare, USA) density gradient centrifugation. To detect surface mobilization of lysosomal associated membrane protein-1 (CD107a), a marker of degranulation associated with cytolytic function, PBMCs were seeded in 96-well plates and cultured alone or stimulated with anti-CD3 antibody (Clone: OKT3, Biolegend) or treated with 50 ng/ml IL-15 for 4 days. 4 hours before the analysis, and anti-human CD107a PE were added to the culture to provoke degranulation. 3 hours before analysis, Brefeldin A were added. Flow cytometry was performed by labeling cells with CD3, CD4, CD28, CD107a expression was quantified by flow cytometry.

### 2.6. Co-culture of CD4^+^CD28^−^ T cells with human renal glomerular endothelial cells

After 24 hours treatment with anti-CD3 antibody and extensive washing, CD4^+^CD28^−^ T cells were co-cultured with human renal glomerular endothelial cells (HRGEC) at a ratio of 1:1 in RPMI 1640 supplemented with 100 mg/ml streptomycin, 100 μg/ml penicillin and 10% fetal bovine serum (FBS) for 48 hours. HREGC were cultured alone or co-cultured with untreated CD4^+^CD28^−^ T cells for 48 hours as a control. Cells from the above co-cultures were washed to remove CD4^+^CD28^−^ T cells and stained the remaining adherent HREGC with Annexin V-FITC and Propidium Iodide (PI) and evaluated for apoptosis by flow cytometry conducting with FCAS Canto II (BD Bioscience).

### 2.7. Multicolour immunofluorescence staining

Tissue samples were obtained from patients with LN. These tissue samples were fixed in formalin, and paraffin embedded prior to sectioning. After deparaffinization, the tissue sections were placed in a plastic container filled with Tris-EDTA buffer for antigen retrieval. All sections were incubated with the primary antibodies to GZMB (496B, eBioscience; dilution 1:200) and CD4 (2H4A2, Protein Tech; dilution 1:400) in humidified boxes overnight at 4°C after blocking with 0.01 M phosphate buffer saline (PBS) containing 3% bovine serum albumin (BSA) for 30 min. Next, the sections were washed and incubated with secondary antibodies (Cy3 conjugated AffiniPure Goat Anti-Rat IgG (GB21302, Servicebio) and FITC conjugated Goat Anti-Mouse IgG (GB22301, Servicebio). The slides were mounted using ProLong Gold antifade reagent with 4’, 6-diamidino-2-phenylindole (DAPI) (G1012, Servicebio) for nuclear staining. Images were acquired with a fluorescence microscope (Leica, Germany).

### 2.8. Cytokine assay

Plasma was separated from fresh EDTA-treated blood samples from SLE patients and healthy controls and stored in frozen. Quantitative analysis for IL-5 was performed using enzyme-linked immunosorbent assay (ELISA) (R&D Systems) according to the manufacturer’s instructions.

### 2.8. Statistics

All quantitative data presented as mean±standard error of the mean (SEM). The significance of the differences for independent samples were determined by unpaired two-tailed Student’s t-test or two-tailed Mann-Whitney U test. The significance of the differences for paired samples were determined by paired two-tailed Student’s t-test or two-tailed Wilcoxon matched-paired signed-rank test. The significance of the differences for samples of more than two group were determined by two-way ANOVA with Tukey’s multiple comparisons or one-way ANOVA with Tukey’s multiple comparisons. P value less than 0.05 was considered to denote a significant difference.

## 3. Results

### 3.1. Single-cell transcriptomic data revealed the existence and characteristics of CD4^+^ cytotoxic T cells

In order to investigate whether CD4^+^ cytotoxic T cells are present in SLE patients, we analyzed scRNA-seq data of 12 SLE and 9 healthy control (HC) samples obtained from the Gene Expression Omnibus (GEO) database (GSE135779, GSE1432016, GSE 157278) (Supplementary Table 2). After quality control, a total of 133,804 cells were obtained for downstream analyses. Thirteen distinctive clusters were identified after PCA and t-SNE analysis and annotated based on the expression of canonical markers. Overall, nine cell types were identified in this step, including T cells, B cells, NK cells, monocytes, macrophages and dendritic cells (Supplementary Fig. 1). We next analyzed the CD4^+^ T cells in T cells, and unsupervised analysis of the transcriptomic data from all CD4^+^ T cells generated a total nine clusters annotated based on the known and high scoring genes (Figs. 1 A and E). We found that the C7 cluster was enriched in genes linked to effector/cytolytic functions such as GZMB, fibroblast growth factor binding protein 2 (FGFBP2), PRF1, granulysin (GNLY), natural killer cell granule protein 7 (NKG7), GZMA, spondin 2 (SPON2), killer cell lectin like receptor G1 (KLRG1), and CC motif chemokine ligand 4 (CCL4) and was annotated as CD4-GZMB cluster (Fig. 1B). We found that the C7-CD4-GZMB cluster was shared among multiple patients and HCs, while the relative proportions of this cluster are different. We also found that the proportion of C7-CD4-GZMB cluster in the CD4^+^ T cells was significantly higher in SLE patients than that in HCs (Fig. 1C). In addition to cytotoxic markers, the C7-CD4-GZMB cluster also highly expressed inflammatory marker and chemotaxis marker (Fig. 1D). Together, these findings indicated that CD4^+^ T cells exhibiting cytotoxic potential and inflammation properties were expanded in patients with SLE.

**Fig. 1.**
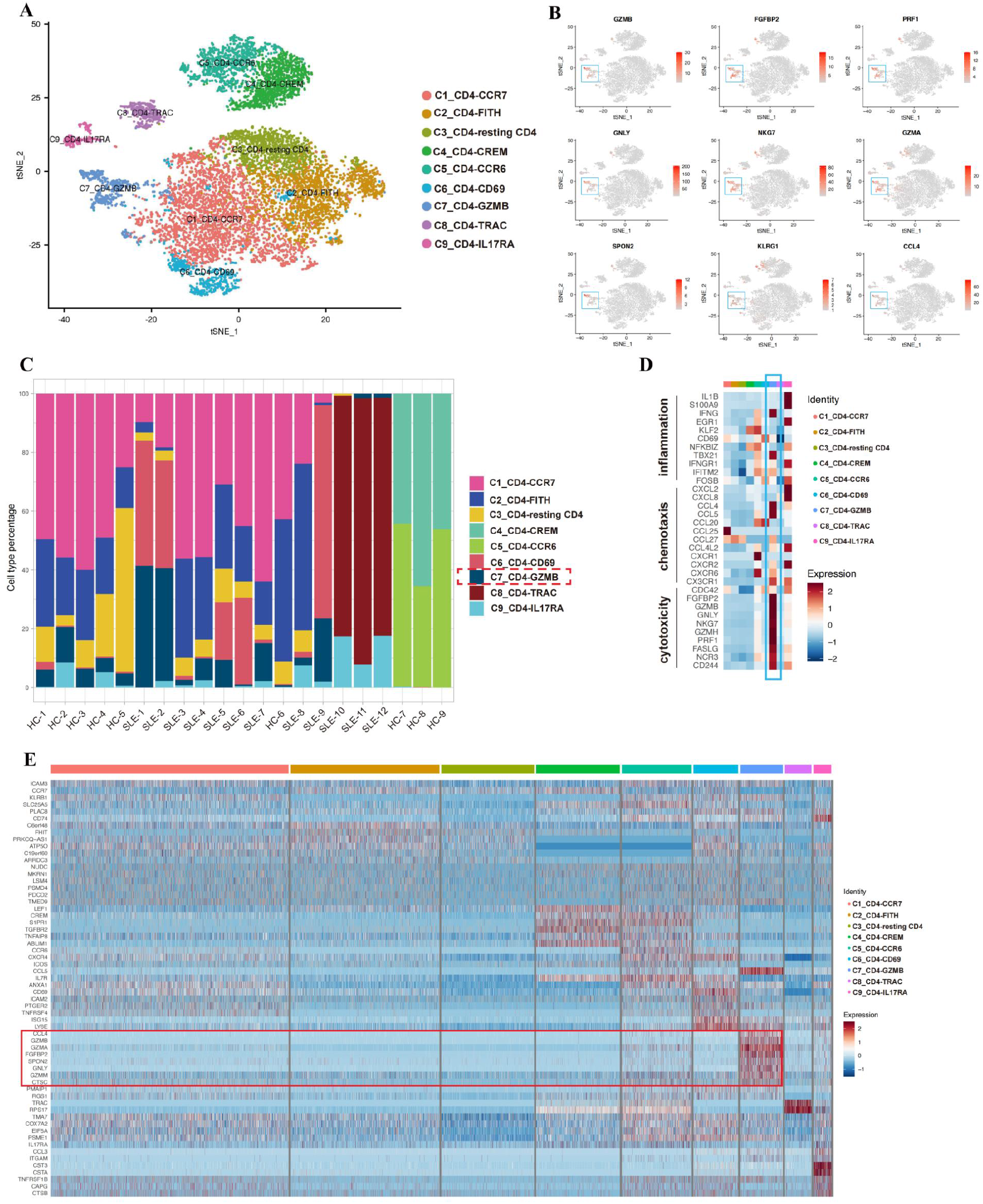
Transcriptomic signature of CD4^+^ cytotoxic T cells from patients with SLE. (A) t-distributed stochastic neighbor embedding (t-SNE) two-dimensional (2D) plot of scRNA-seq data showing the formation of nine clusters. Each dot corresponds to one single cell, with colors corresponding to their respective cell clusters. (B) Expression levels of canonical lineage markers for cytotoxic T cells (C7) across CD4^+^ T cells shown in the t-SNE plots. (C) Relative proportion of nine clusters within the CD4^+^ T cells in SLE patients (n=12) and HCs (n=9). (D) Heatmap showing the average expression levels of genes involved in inflammatory, chemotaxis and cytotoxicity activities of all cells within the nine clusters. (E) Heatmap showing expression of top marker genes of each cell type.

### 3.2. CD4^+^CD28^−^ T cells with effector-memory cytotoxic phenotype abundantly circulate in SLE patients

We next assessed whether CD4^+^ cytotoxic T cells expand in blood samples from SLE patients, using flow cytometry analysis. As shown in Figure 2A and 2C, SLE patients exhibited significantly higher occupancy of CD4^+^CD28^−^ T cells in circulation compared to healthy controls. In addition, SLE patients diagnosed with nephritis showed much higher occupancy than that without nephritis (Figure 2A and C). We found that these cells abundantly expressed cytolytic granule Granzyme B (GZMB; 93%), suggesting that these CD4^+^CD28^−^ T cells function as cytotoxic cells (Figure 2B and C). Furthermore, multicolor immunofluorescence studies showed that in patients with lupus nephritis, CD4^+^GZMB^+^ T cells infiltrated into kidney tissues (Supplementary Figure 2). In addition, a portion of CD4^+^CD28^−^ T cells expressed cytotoxic molecular Granzyme A (GZMA; 20.5%) and Perforin (RPF1; 19.4%) (Figure 2D). The GZMB/PRF1 and GZMB/GZMA staining profiles showed that the PRF1-expressing and GZMA-expressing CD4^+^CD28^−^ T cells were predominantly GZMB positive, suggesting that cytotoxic granules in CD4^+^CD28^−^ T cells were co-expressed (Supplementary Figure 3). In addition to the observed increase in granule molecules, the expression levels of transcription factors Runx3 and T-bet were increased in CD4^+^CD28^−^ T cells (Figure 2F). Furthermore, we found that the chemokine receptor CX3CR1 and NK cell-associated activating receptor NKG2D were also significantly higher in samples from SLE patients compared with those from healthy controls (Figure 2G). Importantly, these cell subsets showed an effector-memory phenotype evidenced by dual loss of CCR7, CD45RA (TEM), together with re-expressed CD45RA (TEMRA) (Figure 2E). Together, these results indicated that CD4^+^CD28^−^ T cells from SLE patients are characterized by both an effector-memory state as well as high cytotoxic potential.

**Figure 2.**
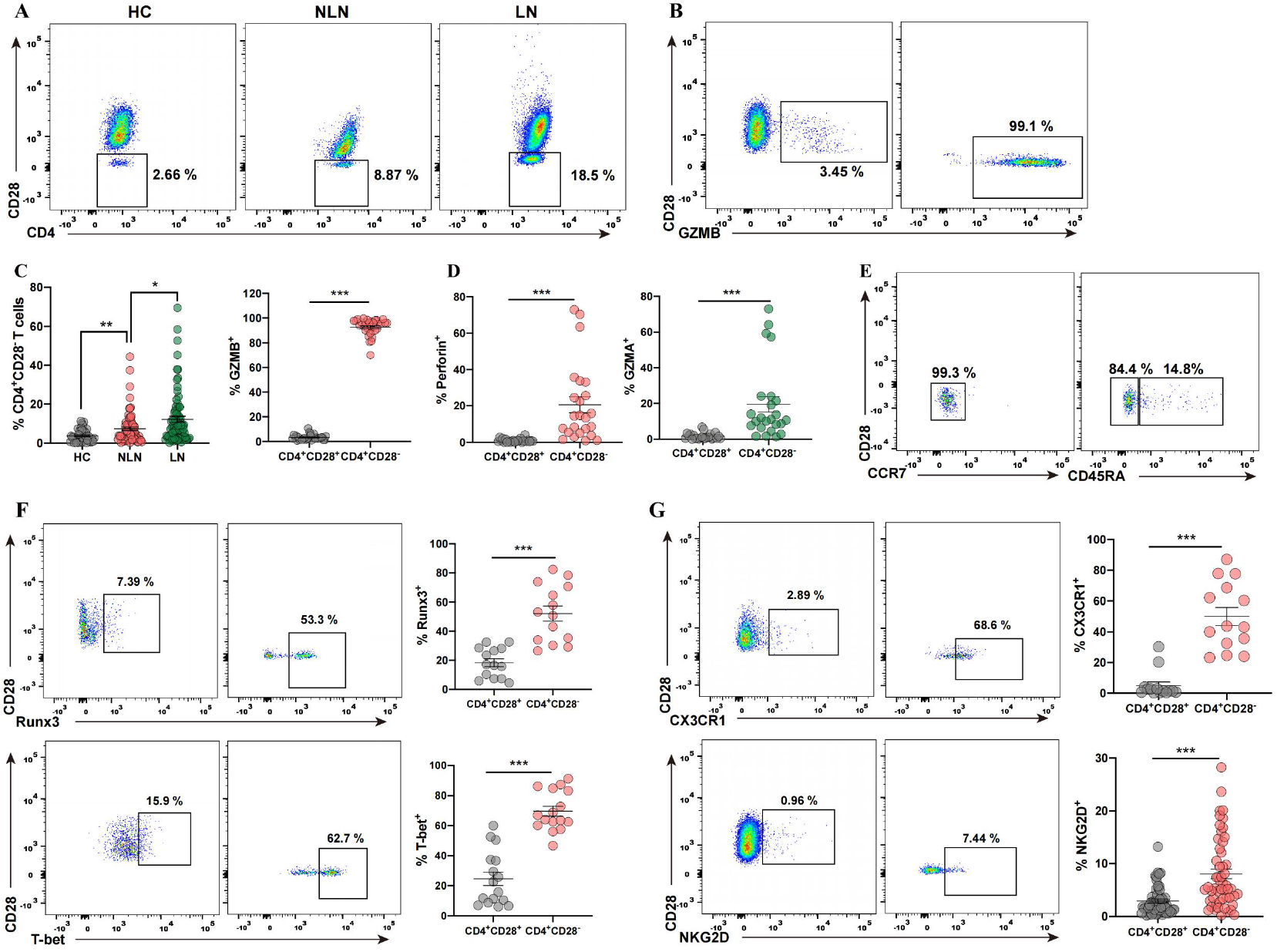
CD4^+^CD28^−^ T cells with effector-memory cytotoxic phenotype are present at high frequency in SLE patients. (A) Frequencies of CD4^+^CD28^−^ T cells in PBMCs from HC, NLN (non-lupus nephritis)-SLE, LN (Lupus nephritis) were determined by flow cytometry. (B) Flow cytometry detection of GZMB expression on CD4^+^CD28^−^ T cells and CD4^+^CD28^+^ T cells from SLE. (C) Graphs showing the percentage of CD4^+^CD28^−^ T cells in HC (n=39), NLN-SLE (n=74), LN (n=80) and the percentage of GZMB^+^ cells in CD4^+^CD28^−^ T cells and CD4^+^CD28^+^ T cells from SLE (n=32). (D) Graphs showing the percentage of GZMA^+^ cells (n=24) and perforin^+^ (n=24) in CD4^+^CD28^−^ T cells and CD4^+^CD28^+^ T cells from SLE. (E) Flow cytometry detection of CCR7 and CD45RA expression on CD4^+^CD28^−^ T cells from SLE. (F) Flow cytometry detection of transcription factor Runx3 and T-bet expression on CD4^+^CD28^−^ T cells and CD4^+^CD28^+^ T cells from SLE; Graphs showing the percentage of Runx3^+^ (n=14) and T-bet^+^ (n=16) cells in CD4^+^CD28^−^ T cells and CD4^+^CD28^+^ T cells from SLE. (G) Flow cytometry detection of CX3CR1 and NKG2D expression on CD4^+^CD28^−^ T cells and CD4^+^CD28^+^ T cells from SLE; Graphs showing the percentage of CX3CR1^+^ (n=14) and NKG2D^+^ (n=53) cells in CD4^+^CD28^−^ T cells and CD4^+^CD28^+^ T cells from SLE. Data information: Data are presented as mean±SEM; \**P* < 0.05, \*\**P* <0.01, \*\*\**P* <0.001, two-tailed Mann-Whitney U test (C, D, G) or unpaired two-tailed Student’s t-test (F).

**Fig. 3.**
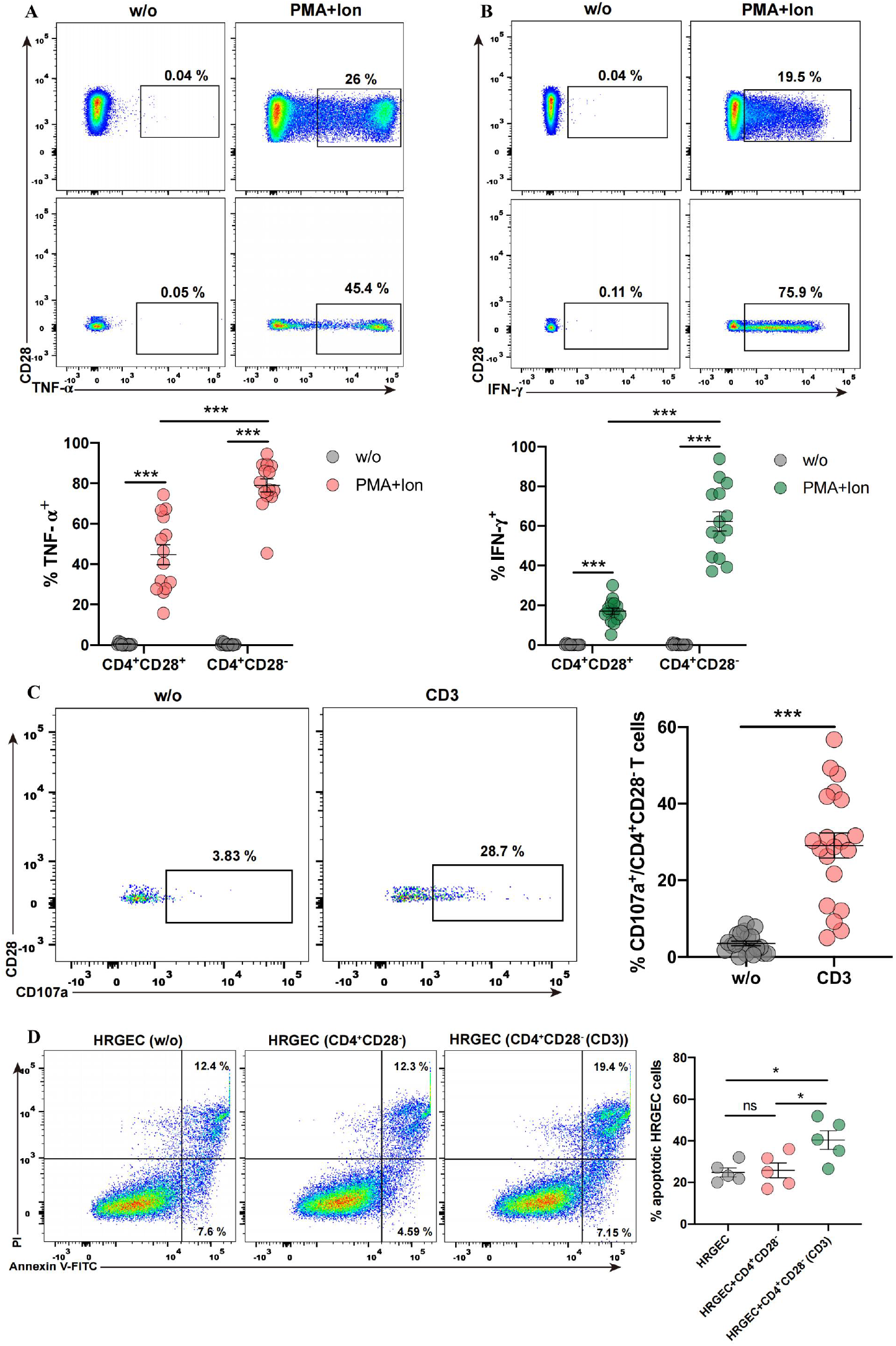
Cytotoxic effect and production of proinflammatory cytokines of CD4^+^CD28^−^ T cells from SLE patients. PBMCs from SLE were cultured alone (w/o) or stimulated with PMA, Ion and Brefeldin A for 4 hours. Percentages of TNF-α (A) and IFN-γ (B) in CD4^+^CD28^−^ T cells and CD4^+^CD28^+^ T cells from SLE (n=14) were measured, respectively. (C) PBMCs from SLE were stimulated with anti-CD3 antibody for 6 hours, and degranulation was quantified with CD107a. Percentage of CD107a in CD4^+^CD28^−^ T cells from SLE (n=20) was determined. (D) HRGECs apoptosis induced by anti-CD3 treated CD4^+^CD28^−^ T cells were detected using flow cytometry. The representative histograms of apoptotic HRGECs in 48h co-cultures, without or with CD4^+^CD28^−^ T cells at a ratio of 1:1 in medium with or without anti-CD3 antibody. Graphs show the percentage of apoptotic HRGECs (n=5). Data information: Data are presented as mean± SEM; \**P* < 0.05, \*\*\**P* <0.001. two-way ANOVA with Tukey’s multiple comparisons (A, B), paired two-tailed Student’s t-test (C), or one-way ANOVA with Tukey’s multiple comparisons (D).

### 3.3. CD4^+^CD28^−^ T cells of SLE patients are functional cytolytic cells that secrete the cytokines TNF-α and IFN-γ

To investigate the functional attributes of CD4^+^CD28^−^ T cells expanded in SLE patients, we measured their inflammatory cytokine production and cytolytic function. As shown in Figs. 3A and B, after stimulating with PMA/ionomycin, both CD4^+^CD28^−^ T cells and CD4^+^CD28^+^ T cells produced more IFN-γ and TNF-α in SLE patients. Moreover, frequencies of IFN-γ and TNF-α producing CD4^+^ CD28^−^ T cells were much higher than that in CD4^+^CD28^+^ T cells, suggesting that CD4^+^CD28^−^ T cells from SLE patients have the potential signature of proinflammatory cytokine. CD107a has been described as a marker of activation-induced degranulation and is a prerequisite for cytolysis. We next assessed the cytotoxicity of CD4^+^CD28^−^ T cells from SLE patients by quantifying CD107a expressed at the cell surface. We found that activation of CD4^+^CD28^−^ T cells with anti-CD3 antibody increased the expression levels of CD107a (Fig. 3C). In addition, treating CD4^+^CD28^−^ T cells with anti-CD3 antibody induced the apoptosis of HRGECs *ex vivo*. Together, these results suggested that cytotoxic CD4^+^CD28^−^ T cells expanded in SLE are functionally active and are capable of producing proinflammatory cytokines.

### 3.4. IL-15 triggers expansion of CD4^+^CD28^−^ T cells in SLE patients

While we successfully validated and characterized CD4^+^ cytotoxic T cells in our study, evidence for their existence remains circumferential, with little direct evidence of how these cells expand or function in the pathogenesis of SLE. Previous studies have shown that the cytokine IL-15 is overproduced in a number of inflammatory and autoimmune diseases. Chronic exposure to IL-15 may promote the expansion of CD4^+^CD28^−^ T cells *in vivo* [39–41]. Here, we investigated whether IL-15 is involved in the expansion and regulate the function of cytotoxic CD4^+^CD28^−^ T cells in SLE. Our results showed that IL-15 was increased in plasma of SLE patients (Supplementary Fig. 5). Furthermore, incubation in IL-15 significantly increased the frequency of CD4^+^CD28^−^ T cells in SLE patients (Figs. 4A and B). Next, we investigated the effects of IL-15 on the proliferation of CD4^+^CD28^−^ T cells in SLE patients by determining the expression of Ki67, a well-known proliferation marker for the evaluation of cell proliferation. The results showed that Ki67-expressed CD4^+^CD28^−^ T cells and the expression level (based on mean fluorescence intensity (MFI)) of Ki67 on CD4^+^CD28^−^ T cells were both significantly increased after stimulation with IL-15. In contrast, minor changes on Ki67-expressed cells and Ki67 expression level were observed in the CD4^+^CD28^+^ T cells followed by IL-15 treatment (Fig. 4C). To identify the active status of CD4^+^CD28^−^ T cells, we measured the expression level of the activation marker CD69. As shown in Fig. 4D, treatment with IL-15 significantly increased the proportion of CD69-expressing CD4^+^CD28^−^ T cells as well as the CD69 expression levels (MFI) on CD4^+^CD28^−^ T cells. We observed only minor differences in the CD69 expression of the CD4^+^CD28^+^ T cells counterparts. Together, these results clearly demonstrated that following IL-15 treatment, CD4^+^CD28^−^ T cells develop a higher capacity of proliferative and higher active status compared with CD4^+^CD28^+^ T cells counterparts. Critically, IL-15 induces the expansion and activation of CD4^+^CD28^−^ T cells, presumably amplifying the pathogenic nature of these cells, thus contributing to the pathogenesis of SLE.

**Fig. 4.**
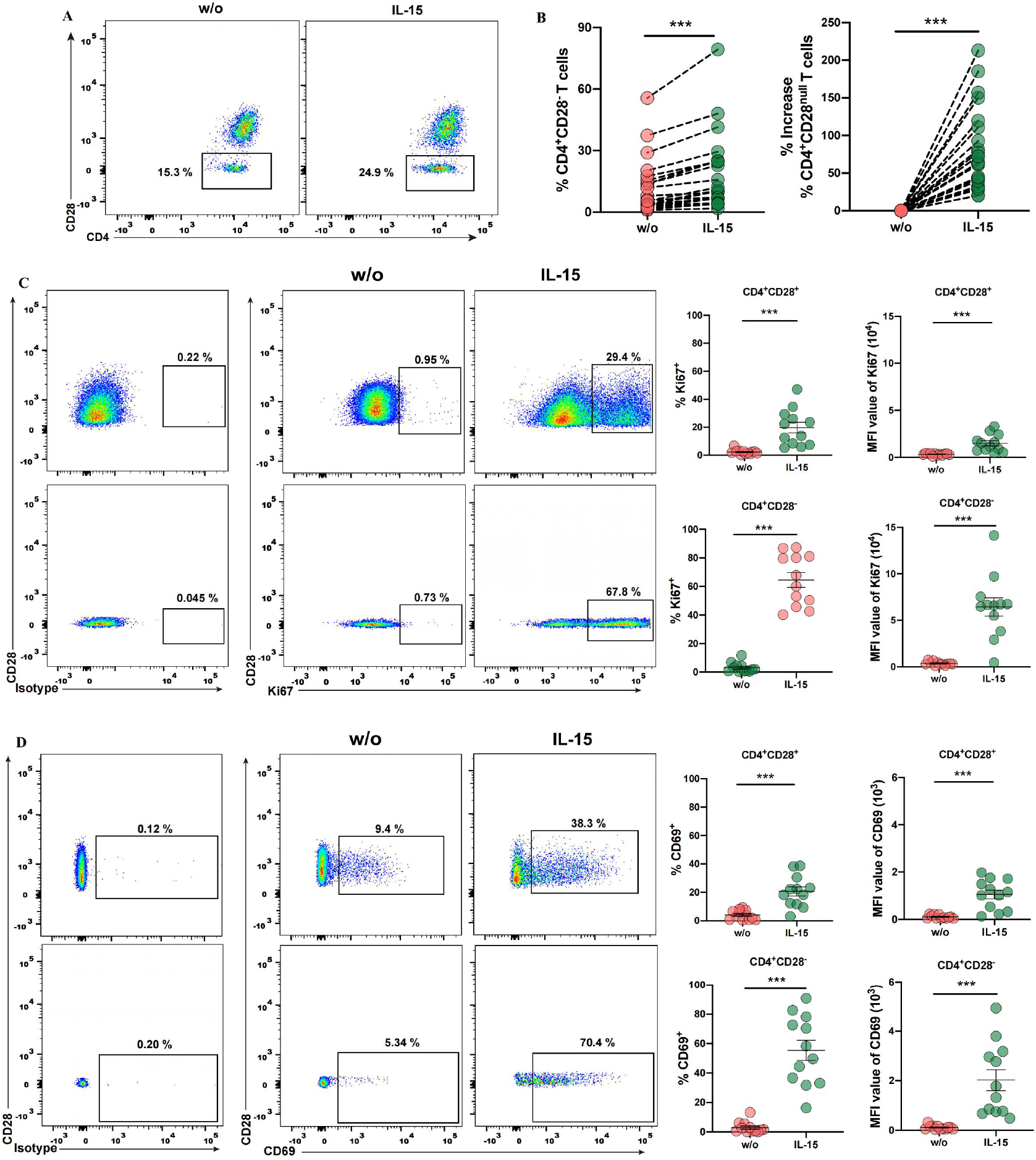
Effects of IL-15 on the expansion of CD4^+^CD28^−^ T cells in Peripheral Blood of SLE patients. PBMCs from SLE were treated with 50 ng/ml IL-15 or without IL-15 (w/o). (A) Representative dot plots of CD4^+^CD28^−^ T cells (n=23) percentage treated with and without IL-15, respectively. (B) Graphs showing the percentage of CD4^+^CD28^−^ T cells (n=23) and % increase of CD4^+^CD28^−^ T cells. (C and D) Representative dot plots of Ki67 expressed and CD69 expressed CD4^+^CD28^−^ T cells and CD4^+^CD28^+^ T cells from SLE, respectively. Graphs showing the percentage of Ki67^+^ (n=12) and CD69^+^ (n=12) cells and the MFI of Ki67 and CD69 in CD4^+^CD28^−^ T cells and CD4^+^CD28^+^ T cells from SLE. Data information: Data are presented as mean±SEM; \*\**P* <0.01, \*\*\**P* <0.001, paired two-tailed Student’s t-test.

**Fig. 5.**
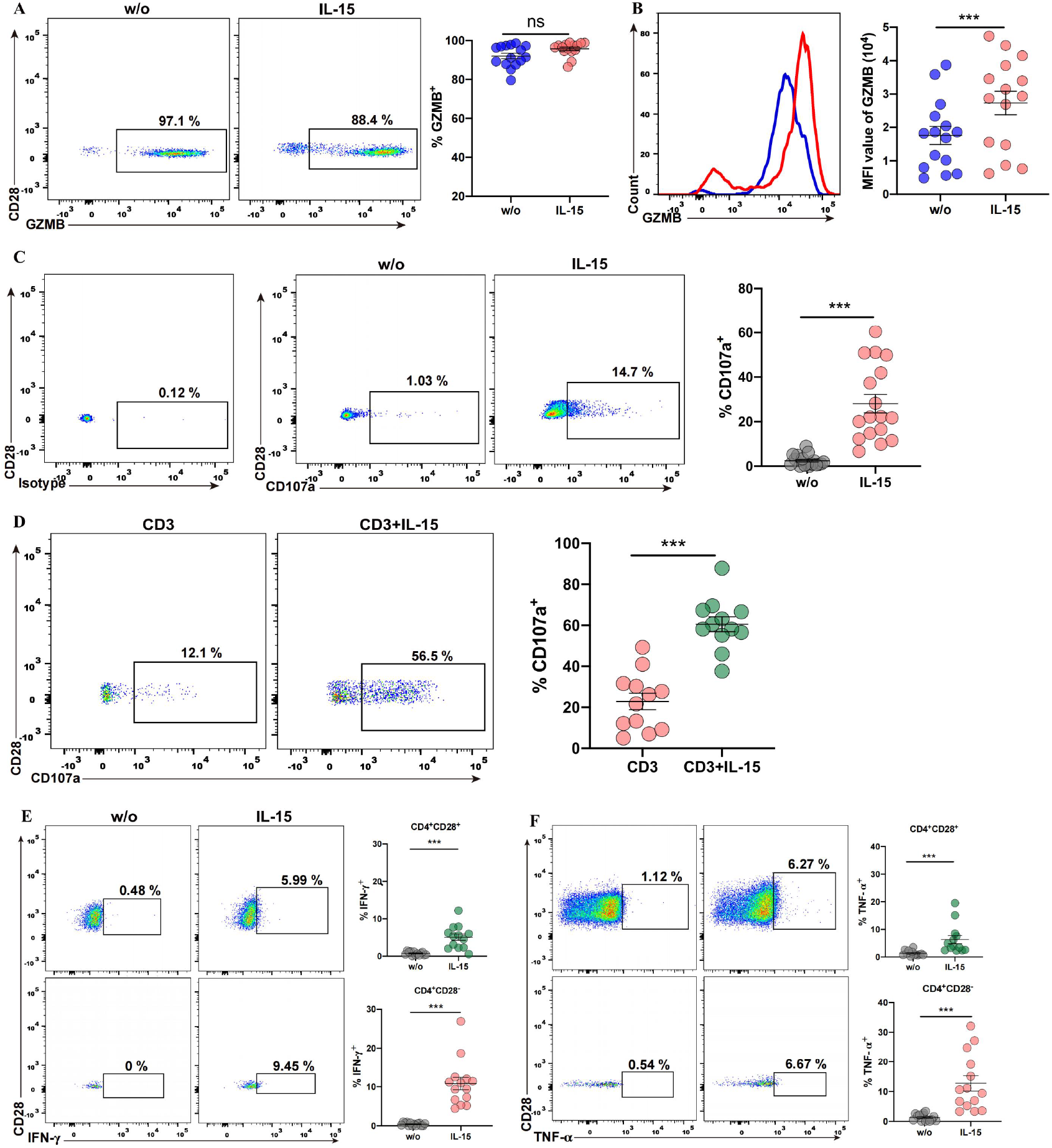
IL-15 increases cytotoxic capacities and cytokines production of CD4^+^CD28^−^ T cells in SLE patients. **(A)** PBMCs from SLE patients were grown in medium alone, with anti-CD3 (1 μg/ml), IL-15 (50 ng/ml), or a combination of both, respectively. (A) Representative dot plots of GZMB expressing on CD4^+^CD28^−^ T cells, graphs showing the percentage of GZMB^+^ cells (n=15) on CD4^+^CD28^−^ T cells when cultured without (w/o) and with IL-15, respectively. (B) Histogram representing the MFI of GZMB on CD4^+^CD28^−^ T cells, graph showing the MFI of GZMB (n=15) on CD4^+^CD28^−^ T cells cultured without (w/o) and with IL-15. (C) Representative dot plots of CD107a expressing on CD4^+^CD28^−^ T cells, graphs showing the percentage of CD107a^+^ cells (n=17) on CD4^+^CD28^−^ T cells cultured without (w/o) and with IL-15. (D) Representative dot plots of CD107a expressing on CD4^+^CD28^−^ T cells, graphs showing the percentage of CD107a^+^ cells (n=12) cultured with anti-CD3 or anti-CD3+IL-15. (E) Representative dot plots of IFN-γ expressing on CD4^+^CD28^+^ and CD4^+^CD28^−^ T cells, graphs show the percentage of IFN-γ^+^ cells (n=14) on CD4^+^CD28^+^ and CD4^+^CD28^−^ T cells cultured without (w/o) and with IL-15. (F) Representative dot plots of TNF-α expressing on CD4^+^CD28^+^ and CD4^+^CD28^−^ T cells, graphs showing the percentage of TNF-α^+^ cells (n=14) on CD4^+^CD28^−^ and CD4^+^CD28^+^ T cells cultured without (w/o) and with IL-15. Data information: Data are presented as mean ± SEM; \*\**P* <0.01,\*\*\**P* <0.001, ns: not significant, paired two-tailed Student’s t-test (A, B, D) or two-tailed Wilcoxon matched-paired signed-rank test (C, E, F).

### 3.5. CD4^+^CD28^−^ T cells from SLE patients exert cytotoxic functions and produce inflammatory cytokine after treated with IL-15

As shown in the previous section, CD4^+^CD28^−^ T cells from SLE patients possess the cytotoxic phenotype with the expression and secretion of cytotoxic granules. Other studies reported that IL-15 induces the expression of cytotoxic effector molecules and promotes the effector capacity of NK and CD8^+^ T cells[42–44]. On the basis of these results, we assessed whether IL-15 treatment affects the expression of granzyme B in CD4^+^CD28^−^ T cells from SLE patients. As shown in Figs. 5A and B, IL-15 did increase the expression levels (MFI) of granzyme B in CD4^+^CD28^−^ T cells, although it did not change the percentage of CD4^+^CD28^−^ T cells expressing granzyme B. Next, we investigated whether the release of cytotoxic molecules was affected by IL-15 using a degranulation assay based on the CD107a expression. PBMCs from SLE patients were treated with IL-15 for 4 days, and the expression levels of CD107a were measured in CD4^+^CD28^−^ T cells using flow cytometry. Our results showed that IL-15 significantly up-regulated the degranulation of CD4^+^CD28^−^ T cells (Fig. 5C). In addition, IL-15 significantly up-regulated the anti-CD3 induced degranulation of CD4^+^CD28^−^ T cells (Fig. 5D). We also investigated whether IL-15 induces the production of disease-related proinflammatory cytokines in CD4^+^CD28^−^ T cells using intracellular flow cytometry. We found that the percentage of cells that produce TNF-α and IFN-γ significantly increased in both CD4^+^CD28^−^ and CD4^+^CD28^+^ T cells treated with IL-15 (Figs. 5E and F), and the increase was higher in CD4^+^CD28^−^ T cells compared to CD4^+^CD28^+^ T cells. Together, our findings strongly suggested that IL-15 stimulates the cytotoxic properties of CD4^+^CD28^−^ T cells, including expression of cytotoxic molecules and an increase in cytotoxic effector function.

### 3.6. IL-15 activates STAT5 and upregulates NKG2D expression in CD4^+^CD28^−^ T cells from SLE patients

Previous studies showed that IL-15 receptor signal is transmitted through the JAK/STAT and RAS/Raf/ERK signaling pathway[45–47]. In order to investigate the mechanism underlying IL-15 regulated function of CD4^+^CD28^−^ T cells, we analyzed the activity of components involved in the IL-15 receptor signaling pathways. As shown in Fig. 6A, we found that STAT5 was phosphorylated following addition of IL-15. In contrast, phosphorylated ERK levels changed little in the same condition (Supplementary Fig. 5). We also found that incubation in IL-15 significantly increased the expression of NKG2D on the surface of CD4^+^CD28^−^ T cells isolated from SLE patients (Fig. 6B). Next, we block IL-15 signaling with the JAK3 selective inhibitor Tofacitinib. As shown in Figure 6C, the expansion and proliferation of CD4^+^CD28^−^ T cells were inhibited after adding Tofacitinib. In addition, our results shown that blocking IL-15 signaling with Tofacitinib also decreased the percentage of Ki67-expressed CD4^+^CD28^−^ T cells and the expression level (MFI) of GZMB in CD4^+^CD28^−^ T cells (Figs. 6D and E). Together, these findings suggested that IL-15 regulate the expansion, proliferation and effector function of CD4^+^CD28^−^ T cells from SLE patients through activation of JAK3/STAT5 and NKG2D signaling pathway.

**Fig. 6.**
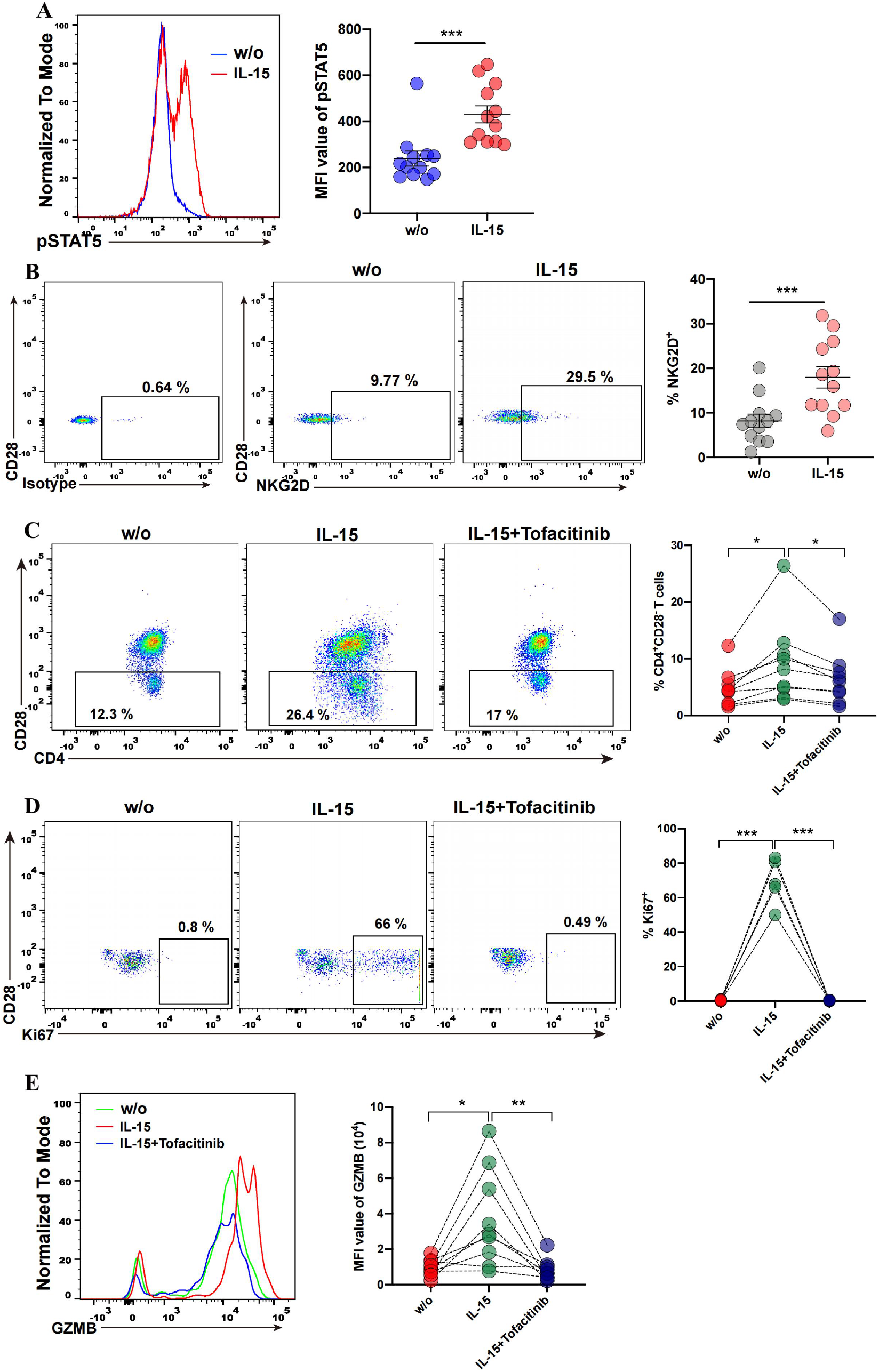
IL-15 activates STAT5 and upregulates the expression of NKG2D in CD4^+^CD28^−^ T cells from SLE patients. **(A)** Phosphorylation levels of STAT5 were determined in CD4^+^CD28^−^ T cells cultured with or without (w/o) IL-15. Histogram and graph showing the phosphorylation levels of STAT5 (n=12). (B) Representative dot plots of NKG2D expressing on CD4^+^CD28^−^ T cells, graphs showing the percentage of NKG2D^+^ cells (n=12) on CD4^+^CD28^−^ T cells cultured without (w/o) and with IL-15. (C) Representative dot plots of CD4^+^CD28^−^ T cells percentage, graphs showing the percentage of CD4^+^CD28^−^ T cells (n=9) untreated (w/o), treated with IL-15 or IL-15 plus Tofacitinib. (D) Representative dot plots of Ki67 expressing on CD4^+^CD28^−^ T cells, graphs showing the percentage of Ki67^+^ (n=5) in CD4^+^CD28^−^ T cells untreated (w/o), treated with IL-15 or IL-15 plus Tofacitinib. (E) Histogram representing the MFI of GZMB on CD4^+^CD28^−^ T cells, graph showing the MFI of GZMB (n=9) on CD4^+^CD28^−^ T cells treated with nothing (w/o), IL-15 and IL-15 plus Tofacitinib. Data information: Data are presented as mean ± SEM; \**P* <0.05, \*\**P* <0.01,\*\*\**P* <0.001, two-tailed Wilcoxon matched-paired signed-rank test (A) or paired two-tailed Student’s t-test (B, C, D, E).

## 4. Discussion

In this study, we systematically analyzed the phenotypic features and cytolytic functions of CD4^+^CD28^−^ T cells in the peripheral blood of SLE patients and identified the effects of IL-15 on the expansion, proliferation and cytotoxic function of CD4^+^CD28^−^ T cells and its mechanism. Our study demonstrated that IL-15 triggers the expansion and cytotoxic function of CD4^+^CD28^−^ T cells, predominantly via the JAK3/STAT5 signaling pathway.

A previous study reported that cytotoxic CD4^+^NKG2D^+^ T cells expand in SLE patients, and showed that this population secretes granzyme and perforin, thus inducing apoptosis of the Treg cells[48]. Here, we demonstrated that the number of cytotoxic CD4^+^CD28^−^ T cells increase in the peripheral blood of SLE patients, especially in patients with lupus nephritis. The percentage of CD4^+^CD28^−^ T cells in SLE patients significantly correlated with SDI score, indicating that CD4^+^CD28^−^ T cells also exert pathogenic function and are involved in SLE-disease tissue damages. In addition, we found that these cells infiltrate the renal tissue of patients with lupus nephritis, where they display a direct cytotoxic activity against HRGECs *ex vivo*. Early studies have shown that the circulating CD4^+^CD28^−^ T cells are considerably expand in patients with Cytomegalovirus (CMV)-seropositive end-stage renal diseases (ESRD), and CMV-associated cytotoxic CD4^+^CD28^−^ T cells potentiate kidney allograft dysfunction by NKG2D-dependent glomerular endothelial injury[49, 50]. In our study, we found that the expression of NKG2D on the surface of cytotoxic CD4^+^CD28^−^ T cells was upregulated in SLE patients. In addition, cell surface expression of chemokine receptor CX3CR1 significantly increased in the CD4^+^CD28^−^ T cells. CX3CR1, the fractalkine receptor, was previously shown to facilitate CD4^+^CD28^−^ T cells’ infiltration in peripheral target tissues[20]. In SLE, a previous study reported that Fractalkine, the ligand for CX3CR1, is upregulated in the serum of patients with SLE[51]. In human proliferation lupus nephritis, the expression of fractalkine and accumulation of CD16^+^ monocytes/macrophages within glomerular were significantly increased. The interaction between fractalkine-expressing cells and CD16^+^ monocytes/macrophages is implicated in pathogenesis of lupus nephritis[52]. Thus, these previous studies established that the high expression of CX3CR1 in CD4^+^CD28^−^ T cells facilitate migration of these cells to renal tissue of lupus nephritis, where they contribute to tissue damage observed in SLE patients. Our analysis found that the expression levels of transcription factors Runx3 and T-bet significantly increased in CD4^+^CD28^−^ T cells. The cooperation of these two factors is thought to activate the cytotoxic program and contribute to the reprogramming of CD4 Th cells into cytolytic effectors[53]. Our results showed that CD4^+^CD28^−^ T cells from SLE patients functioned more as a proinflammatory phenotype than their CD28^+^ counterparts, because the former cell produces higher levels of IFN-γ and TNF-α. Overall, these results suggest that CD4^+^CD28^−^ T cells are characterized by cytotoxic potential and involved in renal tissue injury in SLE patients.

IL-15 is a key manipulator of T-cell function and have been found to promote T-cell survival, proliferation and influence effector functions. The effects of IL-15 on CD4^+^CD28^−^ T cells have been investigated in multiple sclerosis[40], elderly individuals[54], and rheumatoid arthritis[55]. However, whether IL-15 plays a role in SLE is less well established. Previous studies have shown that levels of IL-15 in serum are elevated in SLE patients. This increase has previously been shown to be especially high in patients suffering from lupus nephritis[56, 57]. In agreement with these earlier studies, we found here that the levels of IL-15 in the plasma of SLE patients was higher than that of health controls. Furthermore, we found that cytokine IL-15 play a critical role in regulating the activation and function of CD4^+^CD28^−^ T cells from SLE patients. We dissected the mechanistic basis of IL-15 effects on CD4^+^CD28^−^ T cells from SLE patients and demonstrated that the activation of IL-15/IL-15R/JAK3/STAT5 pathway with the higher levels of phosphorylated STAT5 in CD4^+^CD28^−^ T cells. Tofacitinib, a selective of JAK3 inhibitor that blocks IL-15 signaling, significantly prevents the expansion, proliferation and cytotoxic function of CD4^+^CD28^−^ T cells. Our results suggested that the IL-15/IL-15R/JAK3/STAT5 pathway is an important molecular pathway for IL-15 to regulate the function of CD4^+^CD28^−^ T cells in SLE patients. In addition, our study also found that the expression of NKG2D is upregulated in CD4^+^CD28^−^ T cells followed by IL-15 treatment. An earlier report claimed that the IL-15 receptor signaling pathway couples with NKG2D receptor signaling pathway, thus priming the survival, proliferation and cytotoxic responses of NK cells through JAK3-mediated phosphorylation of DAP10[58]. Another study indicated that IL-15 upregulates the expression and function of NKG2D on CD28-negative effector CD8 CTL activating and inducing the expansion of CD28-negative effector CD8 CTL by arming the NKG2D costimulatory pathway[59]. On the basis of these results, we hypothesize that activated IL-15 receptor cooperates with NKG2D signaling pathway to regulate the function of CD4^+^CD28^−^ T cells. However, to better elucidate the signaling mechanism of IL-15 regulated function of CD4^+^CD28^−^ T cells, further investigation is warranted.

In conclusion, our study demonstrated that CD4^+^CD28^−^ T cells play a critical role in the progression of inflammatory disorder and tissue damage. Importantly, our results suggested that IL-15 contribute to the pathogenesis of SLE by inducing the activation of CD4^+^CD28^−^ T cells via the JAK3/STAT5 and NKG2D/DAP10 pathways. These findings should greatly assist in better understanding the pathological role of CD4^+^CD28^−^ T cells in SLE.

## Acknowledgments

This reaearch was supported by Guangdong Basic and Applied Basic Research Fund (No. 2021A1515111072), the National Natural Science Foundation of China (No. 82201996), the National Natural Science Foundation of China (No. 81971464), and Sanming Project of Medicine in Shenzhen (No. SZSM202111006).

## Appendix A. Supplementary data

**Supplementary data to this article can be found online.**

## Author Contributions

**Xiaoping Hong**: Conceptualization; Resources; Supervision; Investigation; Writing-original draft; Writing-review and editing. **Huan Ren**: Conceptualization; Resources; Supervision; Investigation; Writing-original draft; Writing-review and editing **Tingting Wang**: Investigation; Formal analysis; Writing-original draft; Writing-review and editing; Funding acquisition. **Laiyou Wei**: Investigation; Data curation; Software; Methodology. **Shuihui Meng**: Investigation; Methodology. **Wencong Song**: Resources. **Qianqian Zhao**: Resources. **Yulan Chen**: Formal analysis. **Heng Li**: Investigation; Software. **Zhenyou Jiang**: Supervision; Project administration. **Dongzhou Liu**: Supervision; Funding acquisition.

