## supplementary information for "IL-15 enhances functional properties and responses of cytotoxic CD4^+^CD28^−^ T cells expanded in Systemic lupus erythematosus"

**Supplementary Table-1. Clinical and laboratory features of SLE patients and healthy control**

| **Characteristics** | **NLN-SLE (n=74)** | **LN-SLE (n=80)** | **HC (n=39)** | **P-value** |
| --- | --- | --- | --- | --- |
| Age (mean±SD) | 39.5 ± 13.3 | 37.6 ± 12.1 | 40.8 ± 11.4 | 0.38 |
| Sex (Females) (N (%)) | 67 (90.5) | 73 (91.3) | 37 (94.9) | - |
| Disease duration (year) | 73 (3-120) | 96 (40-138) | NA | 0.014 |
| SLEDAI (median (range)) | 4 (0-16) | 8 (0-20) | NA | <0.001 |
| SDI (median (range)) | 0 (0-2) | 0 (0-3) | NA | 0.06 |
| WBC (×10^9^/l) (median (IQR)) | 4.74 (3.79-6.11) | 5.23 (4.11-7.41) | NA | 0.063 |
| Hemoglobin (g/dl) (mean±SD) | 109.4 ± 23.4 | 107.1 ± 22.8 | NA | 0.45 |
| 24 h proteinuria (g/day) (median (IQR)) | 0.10 (0.08-0.15) | 1.11 (0.22-2.67) | NA | <0.001 |
| Creatinine (median (IQR)) (µmol/L) | 59 (52-67) | 66 (56-96) | NA | 0.001 |
| ESR (mm/h) (median (IQR)) | 19.5 (7-42) | 23 (12-52.5) | NA | 0.14 |
| CRP (mg/L) (median (IQR)) | 1.61 (0.43-3.39) | 1.69 (0.60-4.22) | NA | 0.41 |
| Positive anti-dsDNA (N (%)) | 51 (68.9) | 59 (73.7) | NA | - |
| IgG (g/L) (median (IQR)) | 15.91 (12.29-20.11) | 11.86 (9.06-14.50) | NA | <0.001 |
| IgM (g/L) (median (IQR)) | 0.86 (0.53-1.32) | 0.68 (0.47-1.12) | NA | 0.21 |
| IgA (g/L) (median (IQR)) | 3.00 (2.51-3.59) | 2.37 (1.61-3.02) | NA | <0.001 |
| C3 (g/L) (mean±SD) | 0.70 ± 0.25 | 0.65 ± 0.27 | NA | 0.22 |
| C4 (g/L) (median (IQR)) | 0.12 (0.05-0.17) | 0.12 (0.06-0.19) | NA | 0.54 |

**Notes:** Data expressed as means±SD, medians (interquartile range), or n (%).

**Abbreviation:** N, number in each group; non-lupus nephritis, NLN; lupus nephritis, LN; SLEDAI, systemic lupus erythematosus Disease Activity Index; SDI, systemic lupus international collaborating clinics/ACR damage index; ANA, anti-nuclear Antibodies; Anti-ds-DNA, anti-double-stranded DNA; WBCs, white blood cells; C3 & C4, complement component; ESR, Erythrocyte sedimentation rate; CRP, C-reactive protein.

**Normal levels:** WBCs (×10^9^/l), 4.00-10.0; Hemoglobin (g/dl), 12-18; C3 (mg/dl), 91-185; C4 (mg/dl), 9-63;

**Supplementary Table-2. The information of scRNA- data from GEO database**

| **Name** | **Accession** | |
| --- | --- | --- |
| SLE-1 | aSLE1 | GSE135779 |
| SLE-2 | aSLE2 |  |
| SLE-3 | aSLE3 |  |
| SLE-4 | aSLE4 |  |
| SLE-5 | aSLE5 |  |
| SLE-6 | aSLE6 |  |
| SLE-7 | aSLE7 |  |
| SLE-8 | cSLE10 |  |
| SLE-9 | cSLE14 |  |
| SLE-10 | SLE1 | GSE142016 |
| SLE-11 | SLE2 |  |
| SLE-12 | SLE3 |  |
| HC-1 | aHD1 | GSE135779 |
| HC-2 | aHD3 |  |
| HC-3 | aHD4 |  |
| HC-4 | aHD5 |  |
| HC-5 | aHD6 |  |
| HC-6 | cHD10 |  |
| HC-7 | HC-3 | GSE157278 |
| HC-8 | HC-4 |  |
| HC-9 | HC-5 |  |

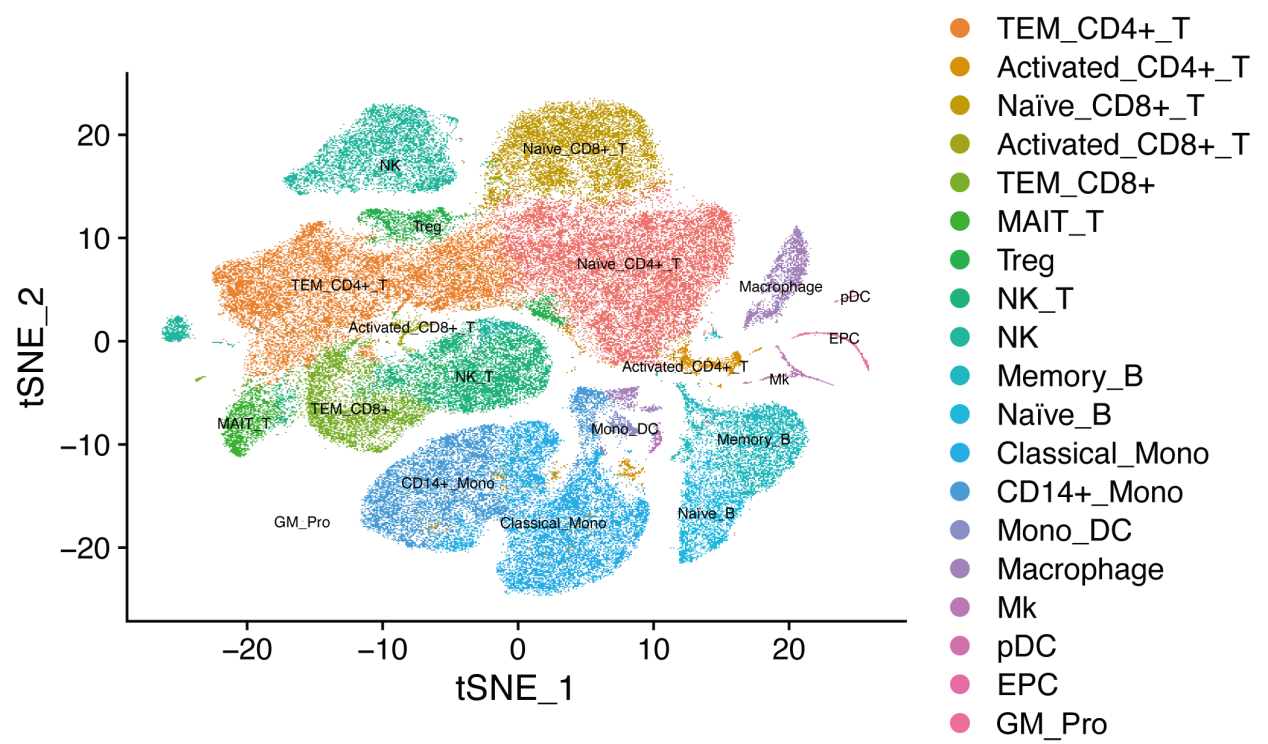

**Supplementary. Fig 1.** T-distributed stochastic neighbor embedding (tSNE) two-dimensional (2D) plot of scRNA-seq data showing the formation of nine clusters. Each dot corresponds to one single cell, colored according to cell cluster.

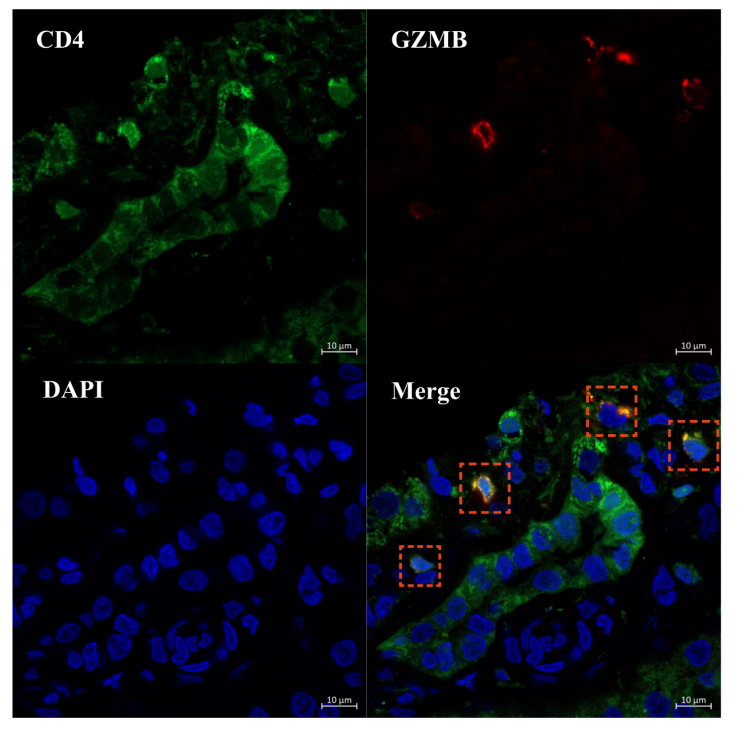

**Supplementary. Fig 2.** Multicolor immunofluorescence image showing CD4 (Green) and GZMB (Red), and DAPI (Bule) staining in a lesion. Red indicate CD4^+^GZMB^+^ T cell.

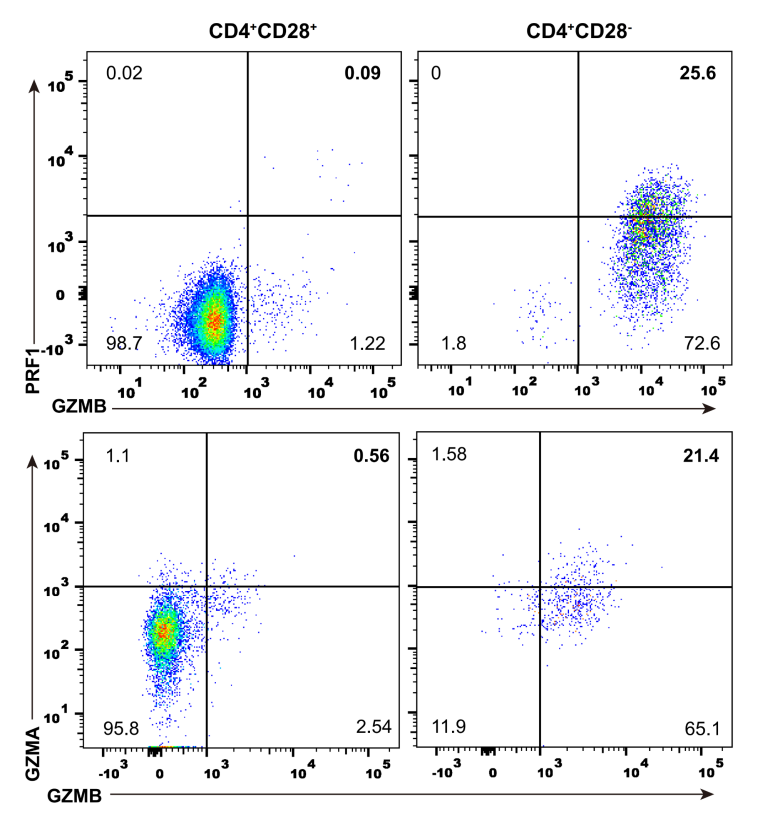

**Supplementary. Fig 3.** The co-expression of two cytotoxic molecules in CD4^+^CD28^-^ T cells and CD4^+^CD28^+^ T cells of SLE were determined with flow cytometry.

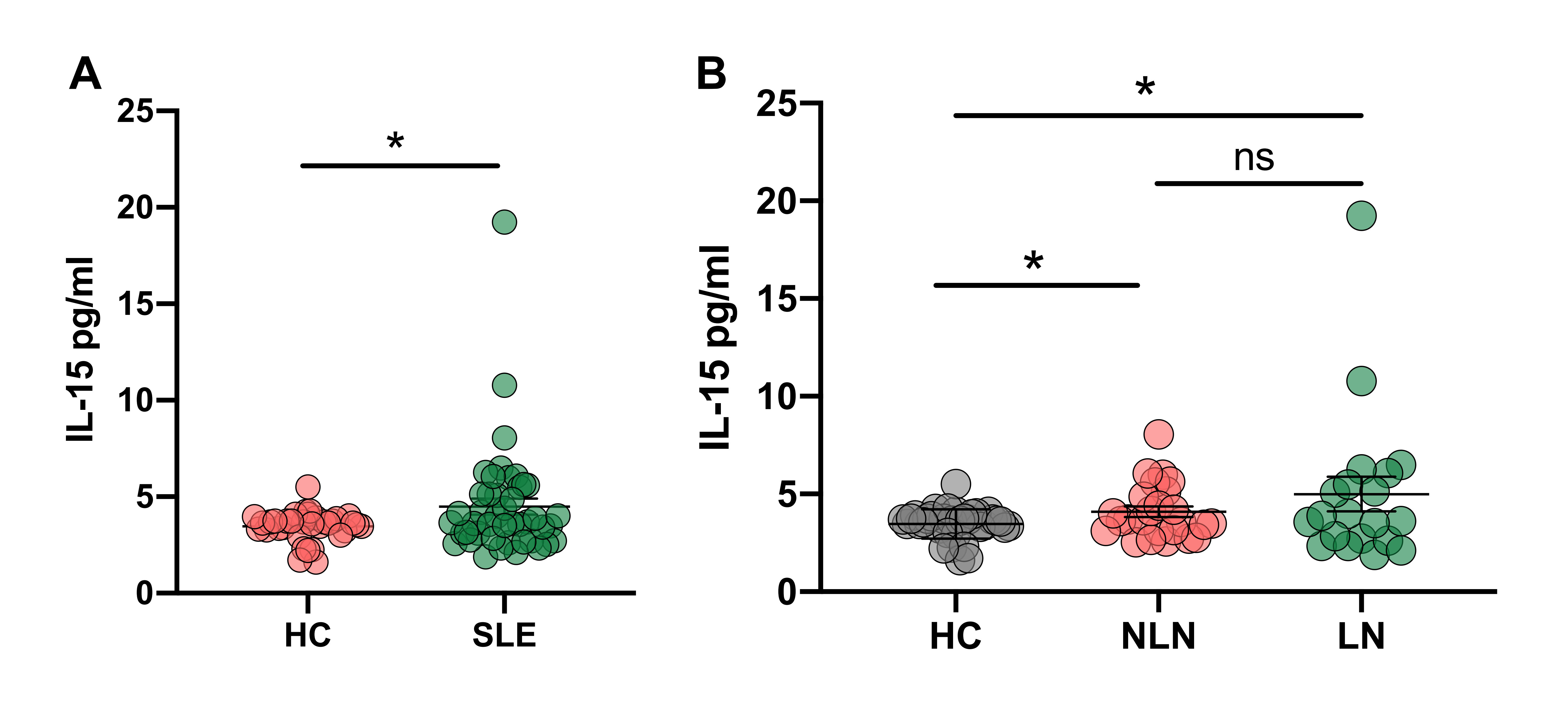

**Supplementary. Fig 4. The cytokine IL-15 levels in serum. ELISA was used to detect the IL-15 in serum samples from the SLE patients and healthy controls.** (A) Graph showing the concentration of soluble IL-15 in SLE patients (n=45) and HCs (n=35). (B) Graph showing the concentration of soluble IL-15 in NLN patients (n=25), LN patients (n=20) and HCs. Data information: Data are presented as mean±SEM; **P* <0.05 , ns: not significant.

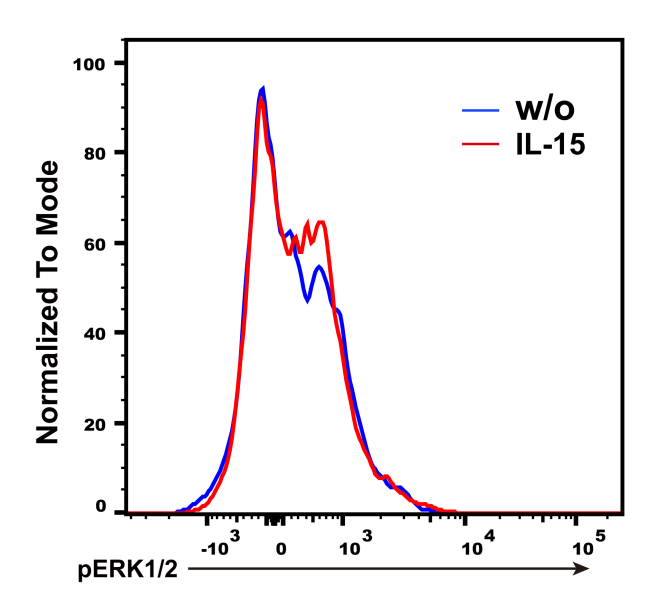

**Supplementary. Fig 5.** The phosphorylation levels of ERK1/2 were determined in CD4^+^CD28^-^ T cells cultured without (w/o) and with IL-15, respectively.
